# Arp2/3 complex activity is necessary for mouse ESC differentiation, times formative pluripotency, and enables lineage specification

**DOI:** 10.1101/2021.11.02.465785

**Authors:** Francesca M. Aloisio, Diane L. Barber

## Abstract

Mouse embryonic stem cells (mESCs), a model for differentiation into primed epiblast-like cells (EpiLCs), have revealed transcriptional and epigenetic control of early embryonic development. The control and significance of morphological changes, however, remain less defined. We show marked changes in morphology and actin architectures during differentiation that depend on Arp2/3 complex but not formin activity. Inhibiting Arp2/3 complex activity pharmacologically or genetically does not block exit from naive pluripotency but attenuates increases in EpiLC markers. We find that inhibiting Arp2/3 complex activity delays formative pluripotency and causes globally defective lineage specification as indicated by RNA-sequencing, with significant effects on TBX3-depedendent transcriptional programs. We also identify two previously unreported indicators of mESC differentiation; MRTF and FHL2, which have inverse Arp2/3 complex-dependent nuclear translocation. Our findings on Arp2/3 complex activity in differentiation and the established role of formins in EMT indicate that these two actin nucleators regulate distinct modes of epithelial plasticity.

**Highlights:**

- Arp2/3 complex activity is necessary for morphology changes during differentiation
- Arp2/3 complex activity regulates transcriptional markers of differentiation
- Inhibiting Arp2/3 complex activity delays entry into formative pluripotency
- Arp2/3 complex activity-dependent shuttling of FHL2 and MRTF occurs in mESCs

## Introduction

As an *in vitro* model, mouse embryonic stem cells (mESCs) derived from the inner cell mass have provided insights on the regulated transition to primed pluripotent epiblast-like cells (EpiLCs) of the post-implantation blastocyst (Martin, 1981; Evans and Kaufman, 1981), which is one of the earliest known transitions in embryonic differentiation (Nichols and Smith, 2009; Nichols and Smith, 2012; Weinberger et al., 2016). While *in vitro* studies with mESCs have revealed how biochemical cues, transcriptional programs, and epigenetics drive differentiation, less is known about morphological changes during differentiation, how they are controlled, and their importance for naive cell differentiation or lineage specification (Gilmour et al., 2017; Villeneuve and Wickström, 2021).

Actin remodeling is a major driver of morphological changes that facilitates diverse cell behaviors (May, 2001; Goley and Welch, 2006). Actin filament architectures are predominantly generated by two classes of actin nucleators: the Arp2/3 complex comprising seven subunits that nucleates branched actin filaments, and formins, a family of fifteen mammalian isoforms that nucleate unbranched actin filaments. While neither class of actin nucleator has been reported for roles in pluripotency transition or lineage specification, processes they regulate, including cellular stiffness (Bongiorno et al., 2018), formation of ventral cortex F-actin asters (Xia et al., 2019), and apparent membrane tension (De Belly et al, 2021), and membrane-to-cortex attachment (Bergert, Lembo et al., 2021), have roles in mESC differentiation. Here we show that mESC morphological changes and actin filament remodeling are dependent on activity of the Arp2/3 complex but not formins, and that Arp2/3 complex activity is necessary for transition from naive mESCs to EpiLCs, including timing entry into intermediate formative pluripotency with global effects on lineage specification.

Formative pluripotency, a recently identified intermediate stage during differentiation of naive mESCs to EpiLCs (Kalkan and Smith, 2014), is considered an executive phase when naive self-renewing signaling networks are dismantled and cells acquire competence for lineage specification. While the formative pluripotent state occurs during spontaneous mESC differentiation, a major limitation for a mechanistic understanding of formative pluripotency timing and regulation is the ambiguity of experimentally isolating and continuously propagating formative pluripotent cells (Smith, 2017; Kalkan et al., 2017; Mulas et al., 2017; Kinoshita and Smith, 2018). Our findings reveal a previously unrecognized role for the Arp2/3 complex in timing entry into formative pluripotency and subsequent lineage specification, which identifies new approaches for studying pluripotency transition states that could be applicable for regenerative medicine. Additionally, our work identifies opposing nuclear localization of myocardin-related transcription factor (MRTF) and FHL2, which are competing co-factors for serum response factor (SRF) transcriptional activity, as previously unreported markers of mESC differentiation and regulated by Arp2/3 complex activity.

## Results

### Inhibiting Arp2/3 complex but not formin activity blocks morphological changes and actin remodeling during mESC differentiation

A mechanism for changes in mESC morphology by actin remodeling and how they are regulated in the context of differentiation and lineage specification remains incompletely understood. Recent advances proposed roles for dynamic cell membrane tension and polarity as requisite regulators of both *in vitro* mESC differentiation (Xia et al., 2019; Bergert, Lembo et al., 2021) and *in vivo* embryonic development (Molé et al., 2021); however, to our knowledge a direct link between actin-dependent changes in morphology and the transcriptional programming of lineage specification has not been reported. To quantify changes in mESC colony morphology in real time we used quantitative DIC imaging of naive E14 mESCs maintained in the presence of LIF2i (Leukemia Inhibitory Factor and pharmacological inhibitors PD0325901 for MEK and CHIR99021 for glycogen synthase kinase-3β) and spontaneously differentiated for 72h after removal of LIF2i (Ying et al., 2008). Naive colonies in LIF2i have a static circular morphology as quantified using *circularity = 4pi(area/perimeter^2)* with a value of 1.0 indicating a perfect circle and values approaching 0.0 indicating an elongated polygon shape (Fig. 1A-B, Supplemental Movie 1). In control cells, colony circularity progressively decreases at 24h, 48h and 72h after removing LIF2i (Fig. 1B, Supplemental Movie 2). Similar morphological changes during mESC differentiation were recently reported (Bongiorno et al., 2018; Bergert, Lembo et al., 2021); however, a regulatory mechanism for dynamic cell and colony morphology was not identified. In determining how these morphological changes are regulated we find that decreases in colony circularity at 48h and 72h-LIF2i are significantly attenuated by CK666, a selective pharmacological inhibitor of Arp2/3 complex activity (Nolen et al., 2009; Yang et al., 2012) but not by SMIFH2, a broad-spectrum inhibitor of formin activity (Rizvi et al., 2009; Ganguly et al., 2015) (Fig. 1A-B). In contrast, we recently showed that SMIFH2 but not CK666 blocks morphological changes during epithelial to mesenchymal transition (EMT) (Rana et al., 2018). Although the selectivity of SMIFH2 was recently shown to also include non-muscle myosins (Nishimura et al., 2021), its ability to inhibit formin activity remains undisputed. Hence, our current findings indicate that the Arp2/3 complex and formins have distinct roles in morphological changes during two different programs of epithelial plasticity, mESC differentiation and EMT, respectively.

**Figure 1.**
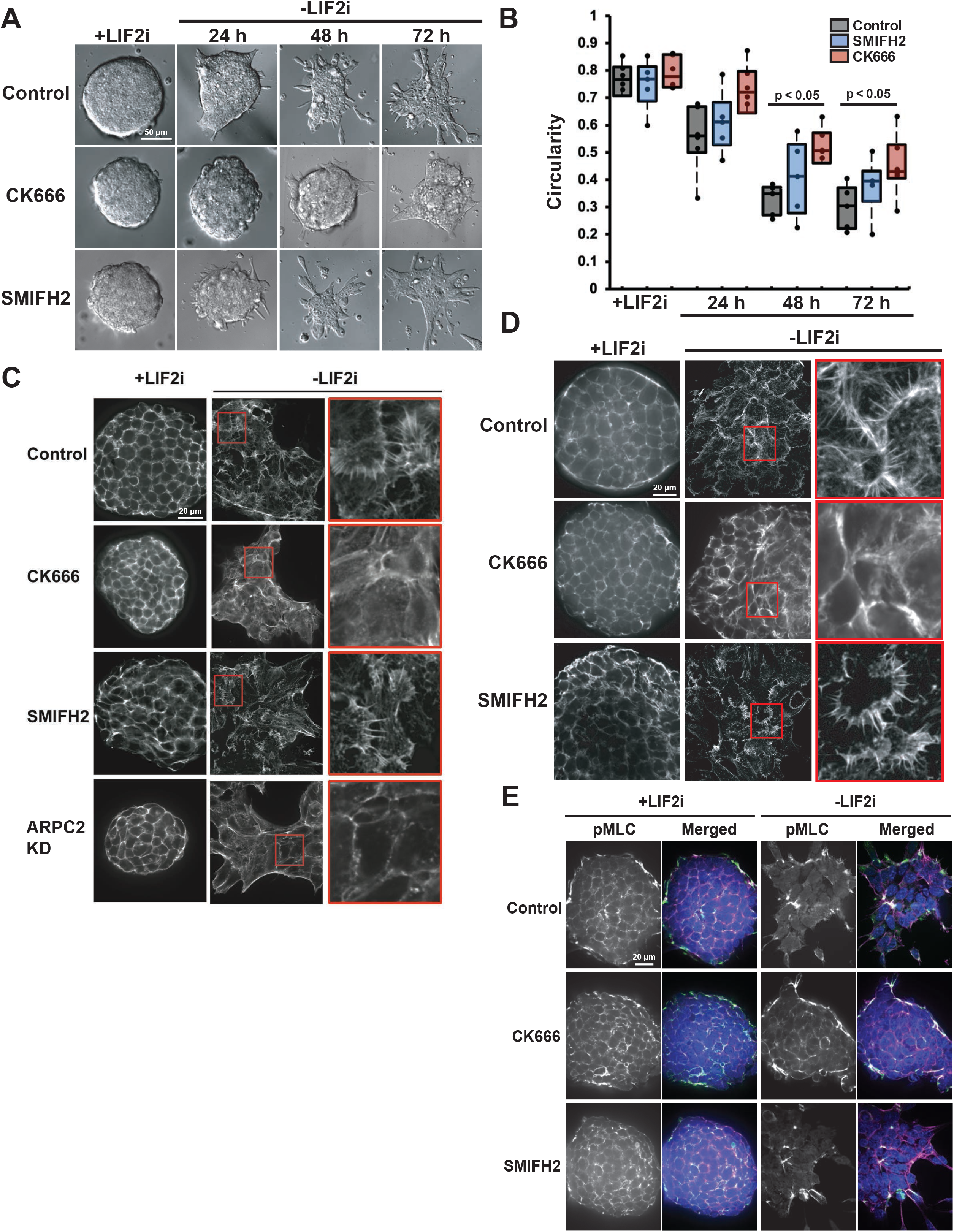
Inhibiting Arp2/3 complex but not formin activity blocks morphological changes and actin remodeling during mESC differentiation. **(A)** DIC images of E14 mESCs over 72h -LIF2i in the absence or presence of CK666 or SMIFH2. **(B)** Colony circularity quantified from DIC images in (A). Box plots show median, first and third quartile, with whiskers extending to observations within 1.5 times the interquartile range, and all individual data points representing means obtained from 6 independent cell preparations of 15-20 individual colonies each **(C)** Confocal images of E14 mESCs +LIF2i and - LIF2i for 72h in the absence or presence of CK666 or SMIFH2 and with Arpc2 KD labeled for F-actin with rhodamine phalloidin. **(D)** Confocal images of V6.5 mESCs +LIF2i and -LIF2i for 72h in the absence or presence of CK666 or SMIFH2 labeled for F-actin with rhodamine phalloidin. **(E)** Confocal images of E14 mESCs +LIF2i and at 72h -LIF2i in the absence or presence of CK666 or SMIFH2 and immunolabeled for pMLC (green) and stained for F-actin with rhodamine phalloidin (magenta) or nuclei with DAPI (blue). Data were analyzed by two tailed unpaired Student’s *t*-test with a significance level of *p*<0.05.

With the established role of Arp2/3 complex in nucleating branched actin filaments, we analyzed actin architectures during differentiation. Using high resolution spinning disc confocal imaging of phalloidin-labeled actin filaments we find that naive E14 mESCs in LIF2i have a compact polygonal cell shape with a cortical ring of actin filaments that are remodeled to an elongated cell shape with prominent membrane protrusions containing ribbed, fan-like actin filaments after 72h -LIF2i (Fig. 1C). In the presence of CK666 but not SMIFH2 actin filaments retain a cortical ring after 72h -LIF2i and fan-like filament networks are rarely seen (Fig. 1C). We also find that effects with CK666 are phenocopied with CRISPR/Cas9 knockdown of ARPC2, an Arp2/3 complex subunit. We confirmed CRISPR/Cas9 editing of the *Arpc2* locus by decreased ARPC2 in E14 mESCs by immunoblotting (Supplemental Fig. 1A-B) and by sequencing (Supplemental Fig. 1E-F). Consistent with the stability of Arp2/3 complex subunits being dependent on their assembly (Di Nardo et al., 2005; Steffen et al., 2006; Rauhala et al., 2013; LeClaire et al., 2015), with *Arpc2* silencing in mESCs there is significantly decreased abundance of the Arp2/3 complex subunit ARP2 (Supplemental Fig. 1C-D). Inhibiting Arp2/3 complex activity in E14 mESCs by either CK666 or Arpc2 knockdown has no effect on the morphology or cortical actin organization of naive cells, but blocks pronounced fan-like actin filaments seen in membrane protrusions after 72h -LIF2i in control E14 cells (Fig. 1C). Similar to our findings with E14 mESCs, we confirmed that genetically distinct V6.5 mESCs show a similar remodeling of actin filament architectures with spontaneous differentiation that is blocked by CK666 but not SMIFH2 (Fig. 1D). Taken together, these findings indicate that actin remodeling with distinct changes in filament architectures occur during mESC differentiation in two mESC lines that are blocked by inhibiting Arp2/3 complex but not formin activity.

Changes in cell morphology are often not driven by actin filament remodeling alone but also in combination with actomyosin contractility (Murrell et al., 2015). In human induced pluripotent cells (iPSCs), a contractile actin fence promotes pluripotency and in mouse embryos suppressing actomyosin contractility regulates epiblast morphogenesis during pre- to post-implantation (Närvä et al., 2017; Molé et al., 2021). Consistent with our finding that CK666 attenuates changes in mESC colony morphology, immunolabeling indicates that phosphorylated MLC (pMLC), an indicator of actomyosin contractility, decorates the cortical actin ring around cells and the peripheral ring around free margins of colonies in control naïve mESCs but is diffuse in the cytoplasm after 72h -LIF2i (Fig. 1E). In contrast, in the presence of CK666 but not SMIFH2 pMLC retains a cortical localization after 72h -LIF2i (Fig. 1E). However, there is no change in pMLC abundance during differentiation in controls or with CK666 or SMIFH2, determined by immunoblotting of E14 mESC lysates (Supplemental Fig. 1G-H). Taken together, these data indicate that activity of the Arp2/3 complex but not formins regulates changes in colony morphology, actin architectures, and localized actomyosin contractility during mESC differentiation.

### Inhibiting Arp2/3 complex but not formin activity impairs differentiation to EpiLCs

We used several approaches to show that Arp2/3 complex activity is also necessary for transcriptional changes during mESC differentiation. We first used a V6.5 dual-reporter (DR) mESC line engineered to express distinct fluorophores as cells transition from naive to primed pluripotency. In brief, work by Parchem et al. (2014) found that V6.5 mESCs in LIF2i express a naive-specific *miR-290* cluster and with spontaneous differentiation upon removal of LIF2i *miR-290* expression decreases and expression of the primed-specific *miR-302* cluster increases. They generated cells that express mCherry driven by the *miR-290* promoter and GFP driven by the *miR-302* promoter. Using flow cytometry, DR mESCs can be used to score for decreased mCherry expression and increased GFP expression as an index of differentiation on the cell population level, while intermediate cells are double-positive for both markers (Fig. 2A). Our analysis indicates that control DR mESCs in LIF2i are >90% mCherry positive (Supplementary Fig. 2A), which is reduced to 16.3% after 72h -LIF2i in controls but is significantly greater at 44.0% with CK666 (Fig. 2B). In contrast, the percent of mCherry single-positive cells is not different with CK689, an inactive analog of CK666 (Nolen et al., 2009), the formin inhibitor SMIFH2, or DMSO as a vehicle compared with controls (Fig. 2B). Further, cell death and proliferation in the presence of CK666, CK689, SMIFH2 or DMSO are not significantly different from control cells at any timepoint during differentiation (Supplementary Fig. 2B, 2E). These data indicate that Arp2/3 complex but not formin activity is necessary for changes in stage-specific miRNA expression during naive to primed pluripotency, suggesting a broader role for Arp2/3 complex-dependent actin remodeling in the context of mESC differentiation beyond changes in morphology.

**Figure 2.**
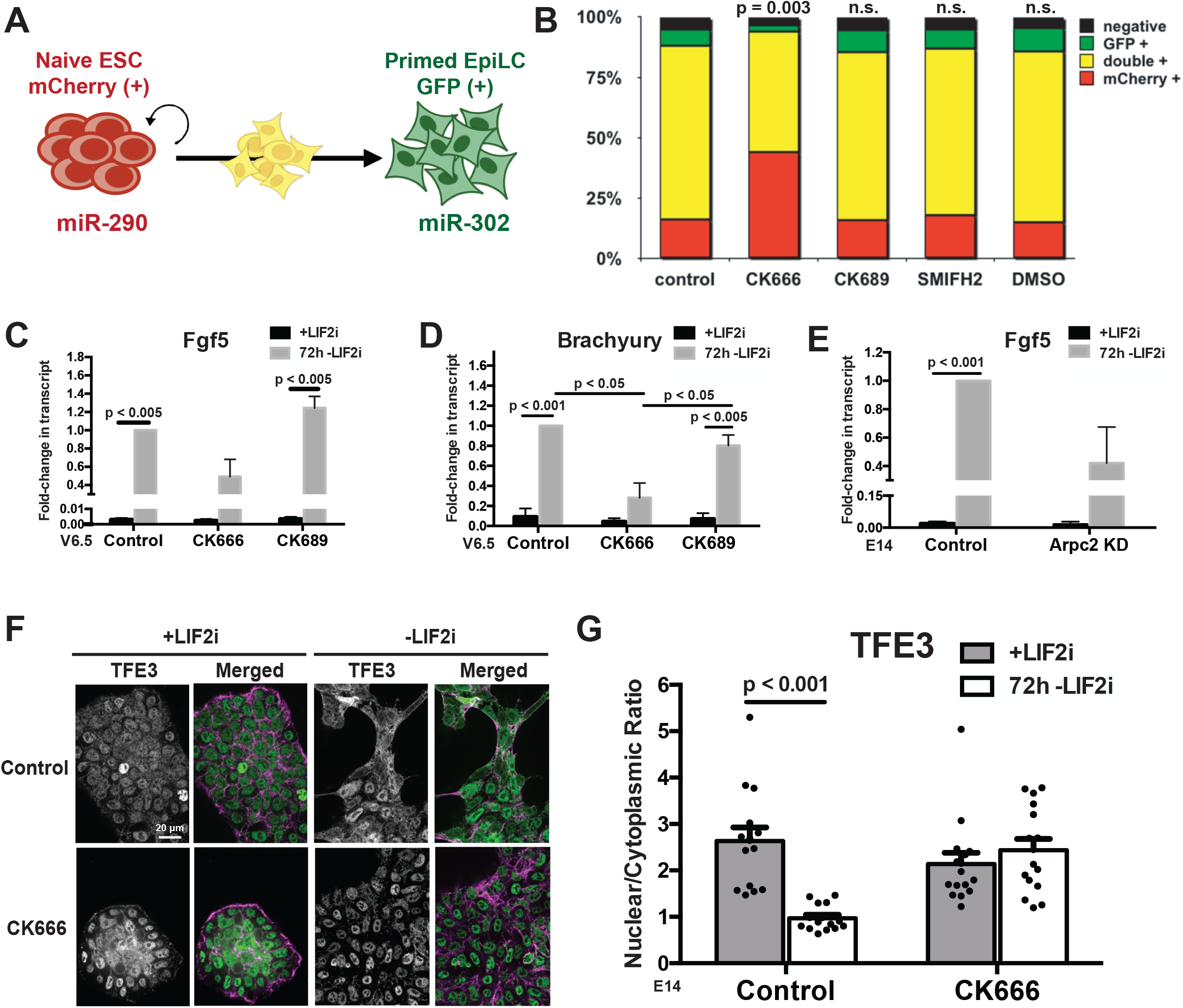
Inhibiting Arp2/3 complex but not formin activity impairs differentiation to EpiLCs. **(A)** Schematic of DR mESCs indicating naive self-renewing mESCs expressing *miR-290-mCherry*, primed EpiLCs expressing *miR-302-eGFP*, and cells transitioning between these stages expressing both markers. **(B)** FACS of V6.5 DR mESCs after 72h -LIF2i in the absence or presence of CK666, CK689 inactive analog of CK666, SMIFH2 or DMSO vehicle, with indicated data representing a mean from 6 independent cell preparations. **(C-E)** RT-qPCR for *Fgf5* **(C)** and *Brachyury* **(D)** in V6.5 DR mESCs and for *Fgf5* in E14 mESCs **(E)** +LIF2i and at 72h -LIF2i. Conditions include the absence (controls) or presence of CK666 or CK689 **(C**,**D)** and control and ARPC2 KD cells **(E)**, with data showing the means ± SEM of 3 independent cell preparations normalized to *TBP*. **(F)** Confocal images of E14 mESCs +LIF2i and at 72h -LIF2i in the absence or presence of CK666 and immunolabeled for TFE3 (green) and stained for F-actin with rhodamine phalloidin (magenta). **(G)** Quantified nuclear to cytoplasmic ratio of TFE3 immunolabeling shown in (F) indicating means ± SEM of 3 independent cell preparations. Data were analyzed by two tailed unpaired Student’s *t*-test with a significance level of *p*<0.05.

As a second approach to test differentiation, we confirmed that CK666 and *Arpc2* silencing attenuates expression of established primed EpiLC markers. RT-qPCR for *Fgf5* (Fig. 2C) and *Brachyury* (Fig. 2D) in V6.5 DR mESCs indicates significantly increased expression in controls and with CK689 after 72h -LIF2i but not with CK666. We used a similar approach to show that expression of *Fgf5* in E14 mESCs significantly increases after 72h -LIF2i in controls but is attenuated with *Arpc2* silencing (Fig. 2E). These data support a role for Arp2/3 complex activity in transcriptional changes associated with differentiation of naive to primed EpiLCs as indicated by pharmacologically or genetically inhibiting Arp2/3 complex activity in V6.5 and E14 mESCs.

Our third approach to test differentiation scored for the cytosolic and nuclear localization TFE3, a bHLH transcription factor that is predominantly nuclear in naive mESCs but mostly cytoplasmic in primed EpiLCs (Betschinger et al., 2013; Villegas et al., 2019; Kalkan et al., 2017). Using quantitative immunolabeling of E14 mESCs, we see that the nuclear to cytoplasmic ratio of endogenous TFE3 significantly decreases after 72h -LIF2i in controls but not in the presence of CK666 (Fig. 2F-G). Taken together, these data reveal a role for Arp2/3 complex activity beyond morphology to include transcriptional indicators such as miRNA expression, primed marker gene expression, and transcription factor localization during transition of naive mESCs to primed EpiLCs.

### Inhibiting Arp2/3 complex activity has no effect on exit from naïve self-renewal but delays entry into formative pluripotency

To further understand how Arp2/3 complex activity enables mESC differentiation, we tested whether it is necessary for exit from naive pluripotency. At 72h -LIF2i, the naive marker *Rex1* (also called *Zfp42*) significantly decreases in V6.5 and E14 cells in the absence and presence of CK666 and CK689 (Fig. 3A-B) as well as in E14 cells with *Arpc2* silencing (Fig. 3B). Moreover, the time-dependent decrease in the expression of *Rex1* as well as *Stra8*, an additional naïve marker, over 120h -LIF2i in E14 cells is similar in the absence or presence of CK666 (Fig. 3C, Supplemental Fig. 2F). These data suggest that although Arp2/3 complex activity is necessary for increased primed EpiLC markers seen in controls, it is not necessary for maintaining naive markers or for exit from naive pluripotency.

**Figure 3.**
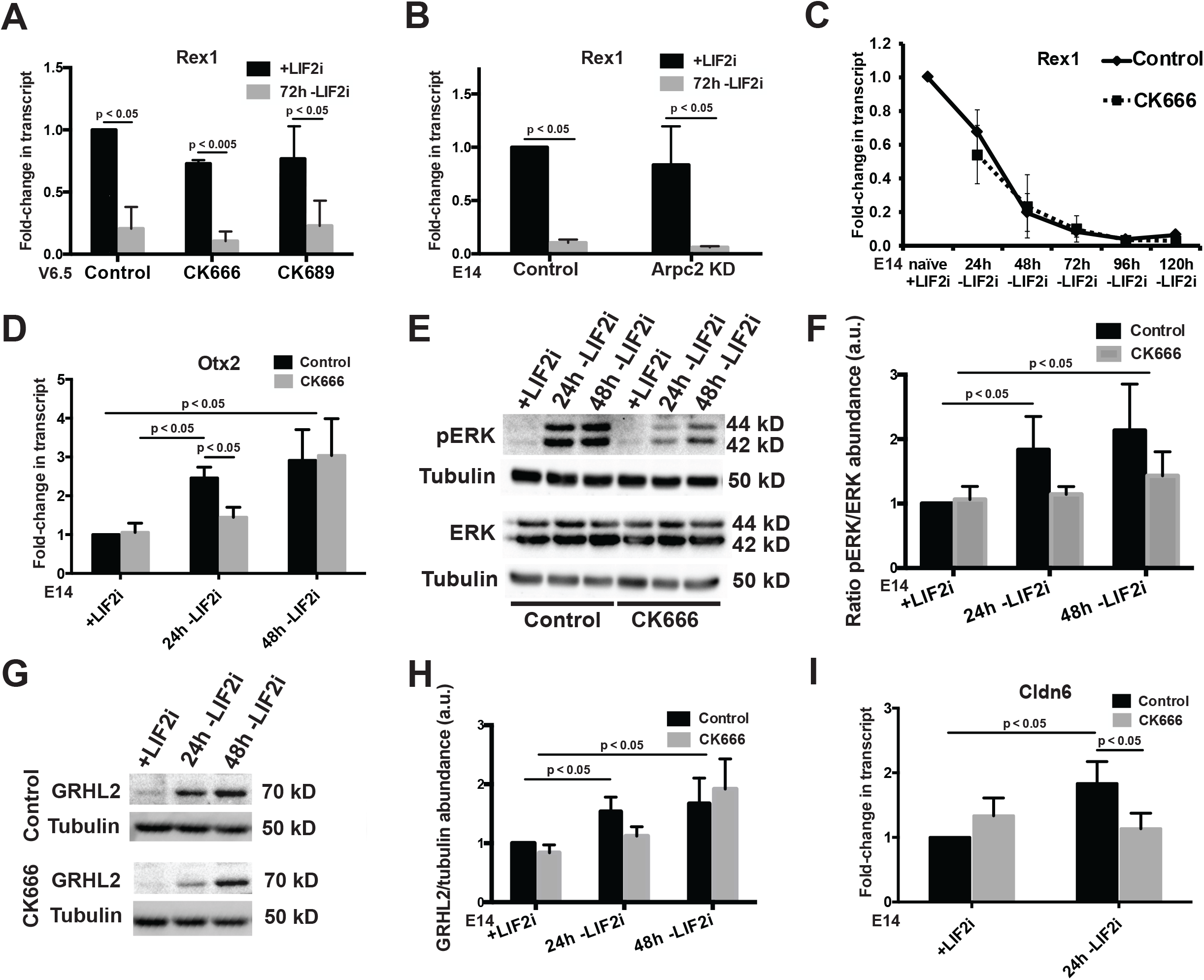
Inhibiting Arp2/3 complex activity has no effect on exit from naive self-renewal but delays entry into formative pluripotency. **(A,B)** RT-qPCR for *Rex1* in V6.5 DR mESCs **(A)** and E14 mESCs **(B)** +LIF2i and at 72h -LiF2i in untreated control cells and in the presence of CK666 or the inactive CK666 analog CK689 **(A)** or in ARPC2 KD cells **(B)**. Data are means ± SEM of 3 independent cell preparations normalized to *TBP*. **(C)** RT-qPCR for *Rex1* in E14 mESCs during 120h time-course -LIF2i in untreated control cells and in the presence of CK666. Data are means ± SEM of 4 independent cell preparations normalized to *TBP*. **(D)** RT-qPCR for *Otx2* in E14 mESCs during 48h time-course -LIF2i in untreated control cells and in the presence of CK666. Data are means ± SEM of 4 independent cell preparations normalized to *TBP*. **(E)** Representative immunoblot of lysates from E14 mESCs during 48h time-course -LIF2i in untreated control cells and in the presence of CK666 probed for pERK, total ERK, or tubulin as a loading control. **(F)** Semiquantitative densitometry of immunoblots described in (E), with data showing means ± SEM of 3 independent cell preparations. **(G)** Representative immunoblot of lysates from E14 mESCs during 48h time-course -LIF2i in untreated control cells and in the presence of CK666 probed for GRHL2 or tubulin as a loading control. **(H)** Semiquantitative densitometry of immunoblots described in (G), with data showing means ± SEM of 7 independent cell preparations. **(I)** RT-qPCR for *Cldn6* in E14 mESCs after 24h -LIF2i in untreated control cells and in the presence of CK666. Data are means ± SEM of 5 independent cell preparations normalized to *TBP*. Given directional *a priori* predictions in panels D-H, data were analyzed by one tailed unpaired Student’s *t*-test with a significance level of *p*<0.05.

An intermediate state between naive and primed pluripotency, termed formative pluripotency, was recently identified. Formative pluripotency is considered an “executive” state when cells are most responsive to differentiation cues and most receptive for lineage commitment (Smith, 2017; Kalkan et al., 2017; Kalkan et al., 2019). Given our findings that inhibiting Arp2/3 complex activity has no effect on exit from naïve pluripotency but attenuates markers of primed pluripotency, we tested whether inhibiting Arp2/3 complex affects expression of intermediate formative pluripotent markers. The formative pluripotent state is currently defined by decreased expression of *Rex1*, which we confirmed is not impaired when Arp2/3 activity is inhibited (Fig. 3A-C), increased *Otx2* (Kalkan et al., 2017; Mulas et al., 2017), increased phosphorylated ERK (pERK) (Kalkan et al., 2019), and increased *Grhl2* (Chen et al., 2018). We confirmed that *Otx2* significantly increases in control E14 cells within 24h -LIF2i (Fig. 3D). In contrast, with CK666 *Otx2* expression at 24h -LIF2i is significantly less compared with control cells and not different than in naive cells (Fig. 3D). After 48h -LIF2i, however, CK666-treated cells have a delayed increase in *Otx2* (Fig. 3D). These results are temporally consistent both with the delayed decrease in colony circularity observed in CK666-treated E14 cells at 48h -LIF2i (Fig. 1A-B) and with previous reports for delayed *Otx2* expression at 48h -LIF2i in the presence of a pharmacological inhibitor of NODAL signaling, which is suggested to function as a timing mechanism for pluripotency transition (Mulas et al., 2017).

Increased pERK, another marker of formative pluripotency, is required for activating downstream formative pluripotent gene regulatory networks (Kalkan et al., 2019; Azami et al., 2019). We find increased pERK in control E14 cells at 24 and 48h -LIF2i compared with total ERK, which does not change during differentiation, as determined by immunoblotting cell lysates (Fig. 3E-F). In contrast, with CK666 pERK does not increase in -LIF2i cells compared with naive cells (Fig. 3E-F). We also used immunoblotting of E14 cell lysates to confirm increased abundance of GRHL2 in control E14 cells at 24 and 48h -LIF2i (Fig. 3G-H), which is similar to reported findings using V6.5 cells (Chen et al., 2018). In contrast, with CK666 increased GRHL2 is delayed with a significant increase at 48h but not at 24h in -LIF2i in cells (Fig. 3G-H). Further, expression of *Cldn6*, a downstream target gene of GRHL2 in mESCs, significantly increases in control E14 cells at 24h -LIF2i but not with CK666 (Fig. 3I). Hence, inhibiting Arp2/3 complex activity in two different mESC lines has no effect on maintenance of naive self-renewal or exit from naive pluripotency but delays entry into the intermediate formative pluripotent stage as indicated by attenuated *Otx2* and *Cldn6* expression as well as pERK and Grhl2 abundance.

### Inhibiting Arp2/3 complex activity disrupts lineage commitment with pronounced effects on TBX3 target genes across all three germ layers

Our findings that Arp2/3 complex activity is necessary for actin remodeling, attenuated expression of primed marker expression, and timing for formative pluripotency during mESC differentiation suggest a role in promoting primed EpiLC lineage specification. To investigate global effects of inhibiting Arp2/3 complex activity on lineage specification, we performed RNA sequencing (RNA-seq) on E14 cells differentiated in the absence and presence of CK666 for 72h -LIF2i and control naive cells maintained in LIF2i (Fig. 4A-B). We found that control naïve +LIF2i and -LIF2i cells have a total of 6,576 differentially expressed genes (DEGs) with an adjusted qval < 0.05 after batch correction (Fig. 4C). Of these DEGs, 1,662 are unique to control -LIF2i cells compared with naïve +LIF2i cells and are not differentially expressed in CK666 -LIF2i compared with naïve +LIF2i cells (Fig. 4C). CK666 -LIF2i cells compared with naïve +LIF2i have 4,796 DEGs, with 457 unique DEGs (Fig. 4C). In CK666 -LIF2i cells compared with both control +/-LIF2i cells, 972 unique DEGs are displayed (Fig. 4A). Gene Ontology (GO) enrichment analysis of this latter subset suggests that unique CK666-specific DEGs are associated with biological processes related to extracellular matrix organization, endothelial cell migration, sprouting angiogenesis, and the MAPK/ERK cascade (Fig. 4D). As a general summary, these data indicate global transcriptomic differences in naive +LIF2i cells, control -LIF2i cells, and CK666-treated -LIF2i cells.

**Figure 4.**
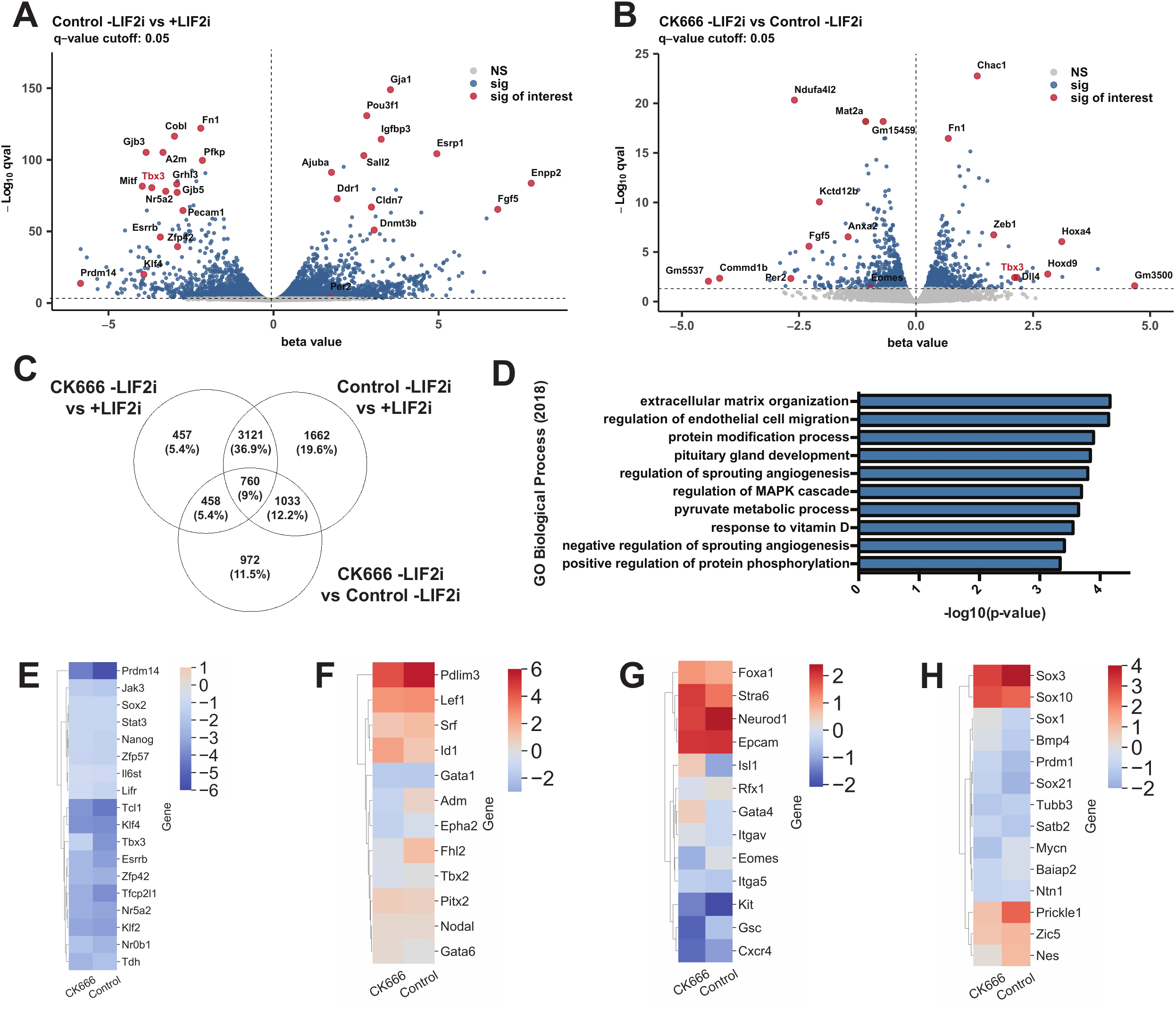
Inhibiting Arp2/3 complex activity causes global defects in lineage specification. **(A)** Volcano plot showing the transcriptome fold-changes (beta values) in Control -LIF2i compared with +LIF2i E14 mESCs after 72h. Each dot represents one gene with significantly changed genes (q-value<0.05) indicated in blue and significantly changed genes of interest indicated in red. **(B)** Volcano plot showing the transcriptome fold-changes (beta values) in CK666 -LIF2i compared with Control -LIF2i E14 mESCs after 72h. Each dot represents one gene with significantly changed genes (q-value<0.05) indicated in blue and significantly changed genes of interest indicated in red. **(C)** Venn diagram showing the number of shared and distinct DEGs indicated by RNA-seq for each listed comparison. **(D)** GO Biological Process (2019) enrichment analysis of 972 DEGs uniquely indicated in CK666 -LIF2i compared to Control -LIF2i after 72h. **(E-H)** Clustermap showing naive mESC marker **(E)**, mesoderm marker **(F)**, endoderm marker **(G)**, and ectoderm marker **(H)** expression indicated by beta values from RNA-seq analysis of E14 mESCs in the absence (Control) or presence of CK666 after 72h - LIF2i compared to +LIF2i.

Consistent with our data indicating that CK666 has no effect on exit from naïve self-renewal (Fig. 3A-C), RNA-seq data also show downregulated naive markers in the presence of CK666 compared with control cells with the exception of *Tbx3* (Fig. 4E). TBX3 is a master regulatory transcription factor known to play dual inhibitory and activating roles as mESCs transition from naive self-renewal to lineage specification (Lu et al., 2011; Kalkan et al., 2019). Consistent with significantly attenuated *Tbx3* expression in CK666 -LIF2i, a number of TBX3 target genes (Russell et al., 2015; Nishiyama et al., 2013; Han et al., 2010) such as *Fgf5, Fn1, Zeb1, Kctd12b*, and *Mat2a* (Fig. 4B) and targets specific to mesoderm such as *Pdlim3, Adm*, and *Fhl2*, (Fig. 4F), endoderm such as *Eomes* and *Kit*, (Fig. 4G), and ectoderm such as *Mycn, Prickle1*, and *Nes* (Fig. 4H) are significantly dysregulated. Taken together, these data indicate a role for Arp2/3 complex activity in timing of formative pluripotent lineage specification related to delayed extinction of naive-promoting targets of *Tbx3*, which is suggested to counteract the initiation of formative pluripotent gene regulatory networks (Kalkan et al., 2019).

TBX3 is an established regulator of early development with dynamic context-dependent roles in embryonic organogenesis across germ layers (Chapman et al., 1996). With its binding to a number of transcription factors such as Klf4, Oct4, Sox2, and Nanog, TBX3 plays a complex role at the center of pluripotency circuitry (Han et al., 2010; Russell et al., 2015) with the potential to act as either an activator or inhibitor of gene expression dependent upon cofactor binding (Carlson et al., 2001). To determine the extent to which inhibiting Arp2/3 complex activity globally affects TBX3 target gene expression, we compared our DEGs to three publicly available mESC datasets related to target genes that change expression relative to a TBX3 reporter (Fig. 5A) (Russell et al., 2015), change expression with shRNA knockdown of *Tbx3* (Fig. 5B) (Nishiyama et al., 2013), and bind TBX3 as indicated by ChIP-sequencing (Fig. 5C) (Han et al., 2010). Comparing data sets shows that CK666 treatment during differentiation generally causes TBX3 target genes to have a contrasting transcriptional profile compared with that of a control differentiation: for each TBX3 gene list, our clustermaps indicate that genes with increasing expression in control -LIF2i had attenuated expression with CK666 and genes with decreasing expression in control -LIF2i had heightened expression with CK666 (Fig. 5A-C). Enriching for TBX3 target genes common to all three datasets that are significantly dysregulated with CK666 compared with control (Fig. 5D) suggests effects on a number of key regulators such *Mycn*, which is essential for neurogenesis (Knoepfler et al., 2002; Kerosuo et al., 2018), *Prdm1*, which has multiple roles in neural fate and germ cell specification (Prajapati et al., 2019; Ohinata et al., 2005), and *Cobl*, which has an actin-related role in neural tube formation (Carroll et al., 2003). Other noteworthy gene expression changes include *Tfe3* (Fig. 2F-G, Fig. 5A) and *Cldn6* (Fig. 3I, Fig. 5A). Additionally, *Eomes*, a TBX3 target gene that plays a context-dependent role in specification of all three germ layers (Costello et al., 2011; Tosic et al., 2019) and *Fhl2*, a mesodermal marker and recently identified tension-dependent actin-binding protein (Sun et al., 2020), are significantly attenuated in the presence of CK666 (Fig. 4F, 5A). Collectively, these data suggest that Arp2/3 complex activity times entry into formative pluripotency, possibly by delayed loss of *Tbx3* expression, resulting in defective downstream global and distinct lineage specification programs.

**Figure 5.**
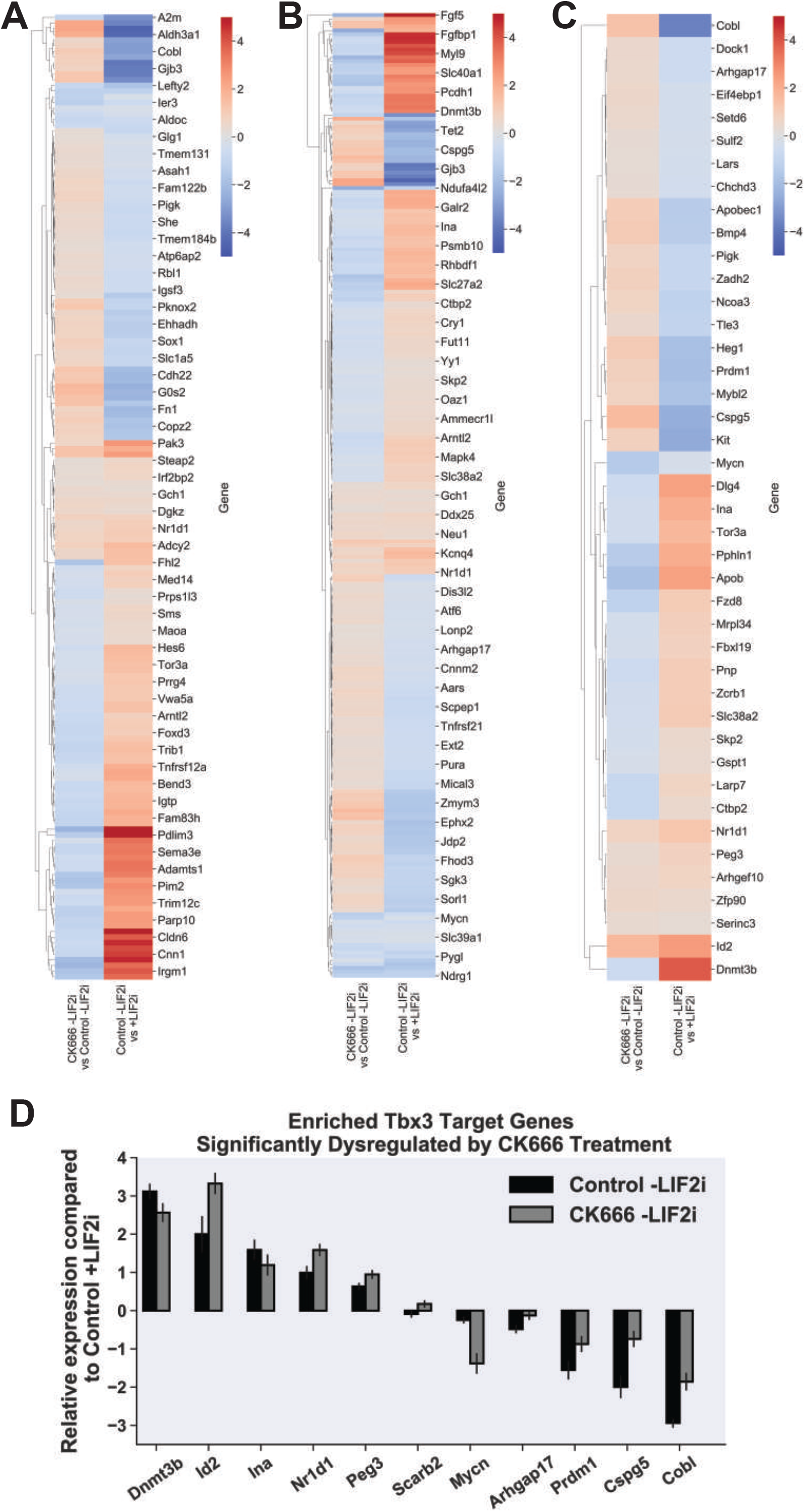
Inhibiting Arp2/3 complex activity disrupts Tbx3-dependent transcriptional programs. **(A-C)** Clustermap showing expression of Tbx3 target genes identified by Russell et al., 2015 **(A)**, Nishiyama et al., 2013 **(B)**, or Han et al., 2010 **(C)** with beta value fold-changes indicated from RNA-seq analysis of E14 mESCs in a control differentiation (Control -LIF2i vs +LIF2i) and how they are affected in the presence of CK666 (CK666 -LIF2i vs Control -LIF2i). **(D)** Enriched bar graph with beta value fold-changes indicated from RNA-seq analysis of E14 mESCs for commonly identified TBX3 target genes across all three published datasets (Russell et al., 2015; Nishiyama et al., 2013; Han et al., 2010) which exhibit significantly different expression (qval < 0.05) in CK666 -LIF2i compared to control -LIF2i.

### Inhibiting Arp2/3 complex activity blocks cytoplasmic and nuclear shuttling of MRTF and FHL2

Our findings on TBX3 target genes regulated by Arp2/3 complex activity led us to identify two previously unreported markers of mESC differentiation – the cytoplasmic and nuclear localization of SRF co-transcriptional activators FHL2 and MRTF. MRTF is an actin polymerization-responsive transcriptional co-activator that, with increased actin polymerization, translocates to the nucleus (Miralles et al., 2003; Posern and Treisman, 2006). FHL2 is a TBX3 target gene and a transcriptional co-activator that is predominantly nuclear in response to decreased F-actin tension (Philippar et al., 2004; Nakazawa et al., 2016). Although formin-dependent nuclear translocation of MRTF is well-described for adult mesenchymal stem cell differentiation, neither MRTF nor FHL2 translocation has been reported to be regulated by Arp2/3 complex activity nor to translocate during mESC differentiation.

We scored for changes in MRTF localization during mESC differentiation and found that in control and SMIFH2-treated naive E14 cells MRTF is diffuse in the cytoplasm but after 72h - LIF2i becomes predominantly nuclear as quantified by a significant increase in the nuclear to cytoplasmic ratio (Fig. 6A, B). In contrast, with CK666 nuclear translocation of MRTF is inhibited with no increase in nuclear abundance at both 72h -LIF2i (Fig. 6A, B) and at 120h -LIF2i (Supplemental Fig. 3A, B). These data indicate that MRTF nuclear translocation occurs during mESC differentiation and is dependent on activity of the Arp2/3 complex but not formins. Further, our RNA-seq data confirm that MRTF target genes (Esnault et al., 2014) have a contrasting transcriptional profile with CK666 compared to controls: our clustermap indicates that MRTF target genes going up in control -LIF2i had attenuated expression with CK666 and genes going down in in control -LIF2i had heightened expression with CK666, including *Fhl2* and *Srf* (Fig. 6C).

**Figure 6.**
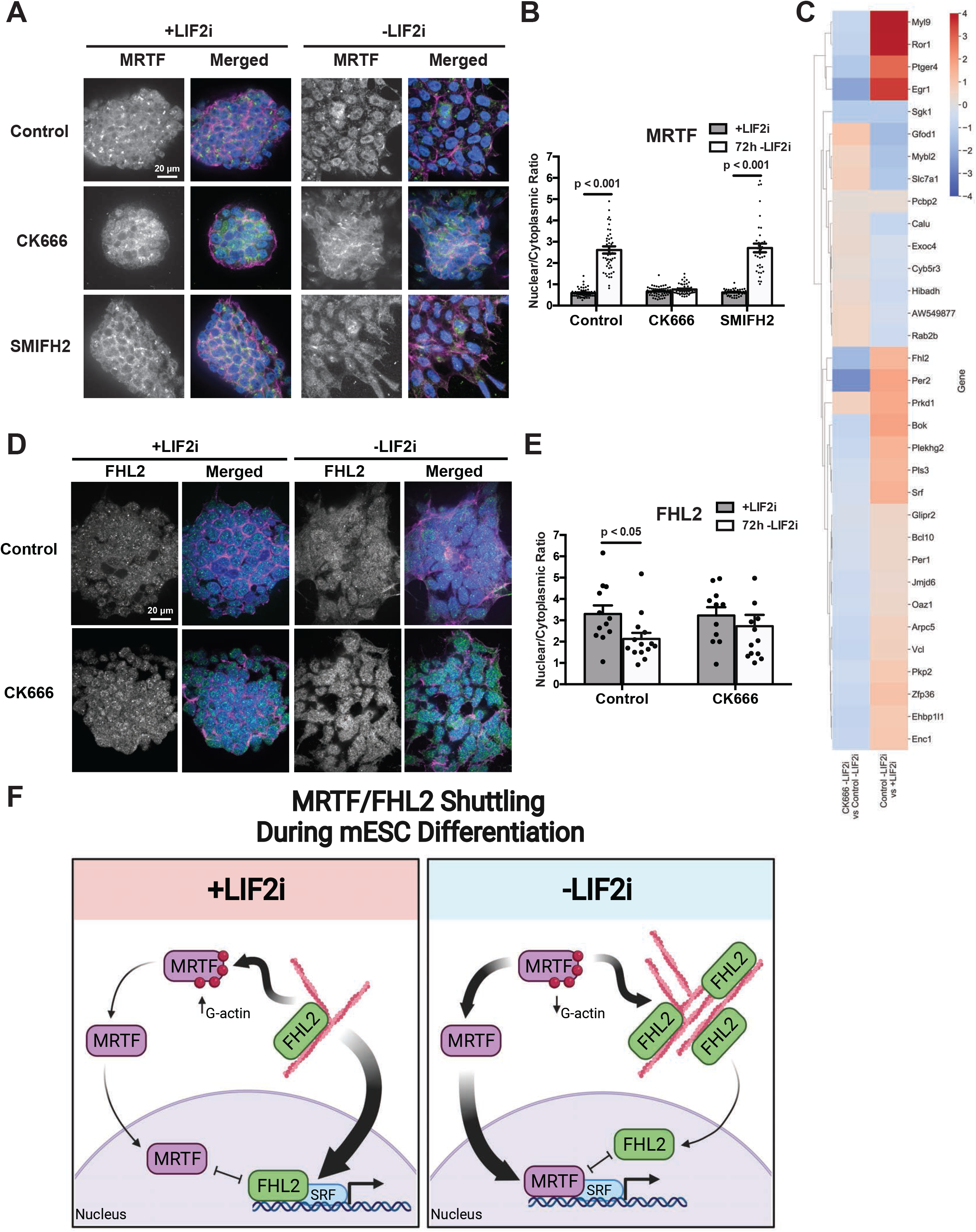
Inhibiting Arp2/3 complex activity blocks cytoplasmic and nuclear shuttling of FHL2 and MRTF. **(A)** Confocal images of E14 mESCs +LIF2i and at 72h -LIF2i in the absence or presence of CK666 or SMIFH2 immunolabeled for MRTF (green) and stained for F-actin with rhodamine phalloidin (magenta) and for nuclei with DAPI (blue). **(B)** Quantified nuclear to cytoplasmic ratio of MRTF immunolabeling shown in (A) indicating means ± SEM of 3 independent cell preparations. **(C)** Clustermap showing expression of MRTF target genes identified by Esnault et al., 2014 with beta values indicated from RNA-seq analysis of E14 mESCs in a control differentiation (Control -LIF2i vs +LIF2i) and how they are affected in the presence of CK666 (CK666 -LIF2i vs Control -LIF2i). **(D)** Confocal images of E14 mESCs +LIF2i and at 72h -LIF2i in the absence or presence of CK666 immunolabeled for FHL2 (green) and stained for F-actin with rhodamine phalloidin (magenta) and for nuclei with DAPI (blue). **(E)** Quantified nuclear to cytoplasmic ratio of FHL2 immunolabeling shown in (D) indicating means ± SEM of cells from images obtained in 4 independent cell preparations. Data were analyzed by two tailed unpaired Student’s *t*-test with a significance level of *p*<0.05. **(F)** Model of competing inverse actin-dependent MRTF/FHL2 nuclear translocation for mESCs in the presence and absence of LIF2i. Created with BioRender.com.

Similar to MRTF, nuclear FHL2 binds with transcription factor SRF to promote target gene expression with noted functions in mesoderm tissues (Lorda-Diez et al., 2018; Renger et al., 2013; Esnault et al., 2014; Philippar et al., 2004; Russell et al., 2015). We scored for changes in FHL2 localization during mESC differentiation and found that in control naive E14 cells FHL2 is nuclear but after 72h -LIF2i undergoes translocation to the cytoplasm, quantified by a significant decrease in the nuclear to cytoplasmic ratio (Fig. 6D, E). In contrast, with CK666 cytoplasmic translocation of FHL2 is attenuated with no significant decrease in nuclear abundance at 72h -LIF2i (Fig. 6D, E). Taken together, these data reveal three previously unrecognized events during mESC differentiation; first is changes in the localization of MRTF and FHL2, second is the opposing nuclear and cytoplasmic translocation of these SRF transcriptional co-activators, and third is that their translocation is dependent on Arp2/3 complex activity (Fig. 6F).

## Discussion

We report a previously unrecognized function of Arp2/3 complex activity in enabling the differentiation of naive to primed mESCs. We find that changes in colony morphology and actin architectures in mESCs are dependent on activity of the Arp2/3 complex but not formins. Our data also indicate that Arp2/3 complex activity is necessary for transition to distinct pluripotency states: although not necessary for exit from the naive state, loss of Arp2/3 complex activity delays entry into the formative pluripotent state, contributing further mechanistic insight on how this newly identified intermediate state is controlled (Smith, 2017; Kalkan et al., 2017; Kalkan et al., 2019). Further, our data include a global examination of actin-dependent lineage specification across all three germ layers in mESCs, which align with evolutionarily conserved roles for force-sensitive differentiation and development in other organisms both and *in vitro* (Chowdhury et al., 2010; Lee et al., 2013) and *in vivo* (Keller et al., 2003; Krieg et al., 2008). Lastly, we show for the first time that MRTF and FHL2, both actin-responsive transcriptional co-activators to SRF, undergo inverse Arp2/3 complex activity-dependent translocation events during mESC differentiation.

We show marked changes in colony morphology and actin architectures during differentiation that depend on Arp2/3 complex but not formin activity. These findings are consistent with reported observations related to morphology and dynamic cellular stiffness during differentiation (Bongiorno et al., 2018) and an acute role for the Arp2/3 complex in mESC actin remodeling (Xia et al., 2019). We observed Arp2/3 complex-dependent actin architectures, which are estabished to generate generate protrusive forces for membrane dynamics (Bailly et al., 2001; Swaney and Li, 2016) and is of particular interest with regard to recently reported roles for dynamic membrane tension related to cortical actin detachment during mESC pluripotency transition (Bergert, Lembo et al., 2021; De Belly et al., 2021). An important question to resolve is how Arp2/3 complex activity regulates transcriptional changes in mESCs compared with its regulation of transcriptional events in other cell models (Yoo et al., 2007; Olson and Nordheim, 2010). Previous studies have indicated force-sensitive lineage specification (Keller et al., 2003; Krieg et al., 2008; Gilmour et al., 2017; Villeneuve and Wickström, 2021) as well as mechanosensing and contractility in fate specification for mESCs (Janmey et al., 2013; Happe and Engler, 2016; Tatapudy et al., 2017). The role of Arp2/3 complex as a central node between biochemical cues and biophysical responses (Iskratsch et al., 2014; Charras and Yap, 2018) suggests a mechanosensitive mechanism whereby Arp2/3 complex activity enables mESC differentiation.

Our data also suggest that Arp2/3 complex activity times entry to intermediate formative pluripotency, an “executive” state when cells are most receptive for lineage specification cues (Smith, 2017; Kalkan et al., 2017; Kalkan et al., 2019). Inhibiting Arp2/3 complex activity delays entry into formative pluripotency, as indicated by the delayed increase in *Otx2* and *Cldn6* expression, and GRHL2 abundance as well as no change in pERK with -LIF2i compared with controls. In related findings, inhibiting NODAL signaling has no effect on exit from naive pluripotency but delays formative pluripotent marker expression from 24h to 48h -LIF2i (Mulas et al., 2017). Taken together, our data indicate that Arp2/3 complex activity is a previously unrecognized node in the growing signaling network of formative pluripotent regulators, which adds mechanistic insight relevant to embryonic lineage specification.

We also reveal that inhibiting Arp2/3 complex activity disrupts lineage commitment across all three germ layers with pronounced effects in TBX3 target genes compared with control cells. TBX3 is a context-dependent master regulator of both naive self-renewal (Niwa et al., 2009; Han et al., 2010; Russell et al., 2015) and lineage specification (Costello et al., 2011; Lu et al., 2011; Weidgang et al., 2013; Kartikasari et al., 2013) and its continued expression during mESC differentiation is reported to destabilize entry into formative pluripotency (Kalkan et al., 2019). Dysregulated *Tbx3* expression is associated with atypical cell and colony morphology in mESCs (Han et al., 2010; Russell et al., 2015), which we also see with loss of Arp2/3 complex activity. Smith and colleagues proposed a relationship between timing of formative pluripotency and RBPJ, a regulator of mESC morphology, whereby RBPJ inhibits TBX3 expression to block formative cells from returning to self-renewal (Kalkan et al., 2019). Further, ERK signaling is reported to inhibit *Tbx3* expression and mESCs null for ERK pathway components have sustained *Tbx3* expression (Niwa et al., 2009; Hamilton et al., 2013; Chen et al., 2015). Our data indicating attenuated pERK and persistent *Tbx3* abundance with CK666 suggest a relationship between Arp2/3 complex activity, mESC morphology, formative pluripotency and TBX3-dependent lineage specification. Future studies on the link between Arp2/3 complex activity and formative pluripotency timing will be important to resolve the interface between morphology and lineage specification, with potential relevance to enhanced protocols for directed differentiation.

We also show that MRTF and FHL2, actin-responsive transcriptional co-activators for SRF, undergo previously unreported translocation events during mESC differentiation. MRTF is a transcriptional co-activator SRF (Posern and Treisman, 2006; Sun et al., 2006; Vartiainen et al., 2007). With increased actin polymerization, MRTF translocates from the cytoplasm to the nucleus where it binds to SRF to promote differentiation programs in adult mesenchymal stem cells (Miralles et al., 2003; Nobusue et al., 2014; McDonald et al., 2015; Bian et al., 2016). Our data show two previously unreported findings on MRTF nuclear translocation; first that it occurs with mESC differentiation and second that it is dependent on Arp2/3 complex activity. Although MRTF has not been reported for direct roles in embryonic differentiation, SRF is confirmed to regulate embryonic mesoderm formation (Weinhold et al., 2000) and *Srf* -/- ESCs have altered cell morphology and reduced cortical actin (Schratt et al., 2002). FHL2, dysregulated in many cancers and developmental disorders, is another SRF-binding transcriptional co-activator which exhibits a direct transcriptional response to the actin cytoskeleton (Sun et al., 2020) and is a TBX3 target gene (Russell et al., 2015). In the cytosol, FHL2 contains LIM domains that mechanoaccumulate on strain sites of tensed actin filaments (Sun et al., 2020). FHL2 is released from F-actin upon loss of filament strain, causing it to translocate to the nucleus where it competes with MRTF for SRF-binding (Philippar et al., 2004). In tandem, MRTF and FHL2 are both direct actin-responsive transcriptional co-activators with dueling roles both in the cytosol and in the nucleus: in the cytosol competing for actin-binding and in the nucleus competing for SRF-binding to promote distinct transcriptional programs. Our data show a previously unreported inverse translocation of MRTF and FHL2 with mESC differentiation. Taken together, these data suggest that inversely mechanosensitive translocation events of MRTF and FHL2 could serve as a novel marker for pluripotency status and provide motivation for understanding how basic cell biology such as actin remodeling can provide a framework for elucidating mechanisms of mESC differentiation and lineage specification.

As recently indicated (Gilmour, 2017; Villeneuve and Wickström, 2021), a current challenge is to identify the connection between the cellular machines that generate shape and the genes that control cell-fate decisions. Our observations, compared with previous findings that activity of formins but not Arp2/3 complex is necessary for EMT and the assembly of unbranched contractile actin filaments (Li et al., 2010; Jurmeister et al., 2012; Rana et al., 2018), indicate that these different classes of actin nucleators and the architectures they generate have selective roles in distinct types of epithelial plasticity. With known functions in migration (Suraneni et al., 2012; Arnold 2008) and adherens junction tension (Verma et al., 2012; Fierro-Gonzales et al., 2012), there are abundant potential mechanisms whereby Arp2/3 complex activity might regulate mESC pluripotency transition (Rotty et al., 2013; Pieters and van Roy, 2014; Wagh et al., 2021; Molé et al., 2021). Our study provides a step toward closing the gap between phenotype and genotype, opening new directions and advancing new approaches to understand how morphological changes and actin filament dynamics promote pluripotency transition, with potential value for additional approaches in directing differentiations for regenerative medicine.

## Materials and Methods

### Cell culture

Wild-type and DR V6.5 ESCs, obtained from R. Blelloch (University of California San Francisco), and E14 ESCs, provided by A. Smith (University of Cambridge) were maintained in tissue culture dishes coated with 0.2% gelatin (G1393; Sigma) at 37°C and 5% CO2 in DMEM (10569; Gibco) supplemented with 15% FBS (FB-11, Omega Scientific, Inc.), glutamine (2 mM), non-essential amino acids (0.1 mM), penicillin-streptomycin (100 U/mL Penicillium and 100 µg/mL Streptomycin), and 2-mercaptoethanol (55 µM). Cells received fresh medium every 24 h and were passaged every three days after dissociating with 0.25% Trypsin-EDTA (25200-056; Gibco). For self-renewal, cells were maintained in medium containing LIF (ESGRO Cat#ESG1106; EMD Millipore) and inhibitors for MEK (1 µM; PD0325901, Cat#S1036; Selleck Chemicals) and glycogen synthase kinase-3β (1 µM; CHIR99021, Cat#S2924; Selleck Chemicals), collectively termed LIF2i. To induce spontaneous differentiation cells were washed in PBS and then incubated in medium without LIF2i as described in Parchem *et al*. (2014) for the indicated times. CK666 (80 µM final; 182515; EMD Millipore), CK689 (80 µM final; 182517; EMD Millipore), and SMIFH2 (25 µM final; S4826; Sigma) were added at 1:000 from stock solutions prepared in DMSO at the indicated times and included in medium replacements every 24h. Mycoplasma contamination was tested 2x/year using media obtained from cells maintained for 48h in the absence of penicillin-streptomycin by using a PCR Mycoplasma Detection Kit (abm Cat#G238). Cell lines were authenticated commercially by IDEXX BioAnalytics (USA).

### CRISPR/Cas9 gene editing

The validated guide RNAs (gRNA) targeting the Arpc2 locus were selected from the Genome-scale CRISPR Knock-Out (GeCKO) v2 mouse library (www.genome-engineering.org) (Sanjana et al., 2014). After annealing and adding BbsI cut site overhangs, candidate gRNAs were cloned into the pSpCas9(BB)-2A-GFP (PX458) plasmid vector (Addgene plasmid #48138; RRID: Addgene_48138) (Ran et al., 2013). At 48 h after transfecting cells with plasmids, single GFP(+) cells were sorted by fluorescence-activated cell sorting as described below. Edited clones were validated by PCR and sequencing (forward primer AGCTGTTGAATGCAATGAGG, reverse primer TCCTCTGGGTAAAGGACCT) and confirmed by immunoblotting as described below. TIDE webtool (https://tide.deskgen.com) was used to quantify editing efficacy and identify the predominant type of indel in the edited clone (Brinkman et al., 2014). The gRNA sequence used to generate the confirmed Arpc2 edited clone was as follows: TTCTTGGTAAATCCAGAACC.

### DIC image acquisition and quantitative analysis

For DIC imaging, naïve E14 ESCs were plated for 24h on gelatin-coated glass bottom microwell dishes (P35G-1.5-14-C; MatTek) in medium containing LIF2i, washed with PBS, and then maintained for the indicated times in medium without LIF2i. CK666 and SMIFH2, as indicated above, were added at the time of LIF2i removal and replaced every 24h until completion of imaging. Live cells were imaged using a Plan Apo 40 0.95 NA objective on an inverted spinning disc microscope system (Nikon Eclipse TE2000 Perfect Focus System; Nikon Instruments; Nikon Instruments) equipped with D-C DIC Slider 40x I (MBH76240; Technical Instruments), a multipoint stage (MS-2000; Applied Scientific Instruments), a CoolSnap HQ2 cooled charge-coupled camera (Photometrics) and camera-triggered electronic shutters controlled with NIS-Elements Imaging Software (Nikon). Approximately 15-20 colonies were imaged for each condition and time point. Colony circularity was quantified using the ImageJ plug-in “Circularity” feature. In brief, this feature is an extended version of the Measure command in ImageJ that calculates object circularity using the formula *circularity = 4pi(area/perimeter^2)*, with a circularity value of 1.0 indicating a perfect circle. As the value approaches 0.0, it indicates an increasingly elongated polygon. Statistical analysis was performed with GraphPad Prism 6 software.

### Immunolabeling, staining, and image acquisition

For immunolabeling, cells were plated on gelatin-coated coverslips prepared in an ultrasonic cleaning bath. In brief, coverslips were sonicated for 20 minutes in the presence of ddH2O and Versa detergent, washed in ddH2O, sonicated again for 20 minutes, and stored in 70% EtOH. Cells were maintained for the indicated times, washed with PBS, and fixed with 4% formaldehyde for 15 min at RT. Cells were then permeabilized with 0.1% Triton X-100 for 5 min, incubated with blocking buffer of 5% horse serum and 1% BSA in PBS for 1h, and then incubated with primary antibodies overnight at 4°C. The cells were then washed with PBS, incubated for 1h at RT with secondary antibodies conjugated with fluorophores, and washed with PBS. One wash included Hoechst 33342 (1:10,000; H-3570; Molecular Probes) to stain nuclei. Primary antibodies included Phospho-Myosin Light Chain 2 Thr18/Ser19 E2J8F (1:200; #95777; Cell Signaling Technology), MRTF-A-C19 (1:200; sc-21558; Santa Cruz Biotechnologies), FHL2 (1:200; HPA006028; Sigma), and TFE3 (1:200; 14480-1-AP; Proteintech). Actin filaments were labeled with rhodamine-phalloidin (1:400; Invitrogen) added during secondary antibody incubations. Cells were imaged using a 60X Plan Apochromat TIRF 1.45 NA oil immersion objective on an inverted microscope system (Nikon Eclipse TE2000 Perfect Focus System; Nikon Instruments) equipped with a spinning-disk confocal scanner unit (CSU10; Yokogawa), a 488-nm solid-state laser (LMM5; Spectral Applied Research), a multipoint stage (MS-2000; Applied Scientific Instruments), a CoolSnap HQ2 cooled charge-coupled camera (Photometrics) and camera-triggered electronic shutters controlled with NIS-Elements Imaging Software (Nikon). Nuclear-to-cytoplasmic ratios were quantified using NIS-Elements Imaging Software (Nikon). Briefly, the fluorescence in the nucleus and cytoplasm were manually sampled by selection of regions-of-interest either colocalized with nuclear DAPI or not. The ratio of fluorescence was then calculated by dividing the nuclear fluorescence intensity with that of the cytoplasm for a given cell. Statistical analysis was performed in Excel (Microsoft) using two-tailed t-test.

### Flow Cytometry

DR ESCs and CRISPR-Cas9 edited E14 ESCs were prepared for flow cytometry by washing with PBS at the indicated times, dissociated with 0.25% Trypsin-EDTA, and collected by centrifuging at 1000 rpm for 3 min at room temperature. Pelleted cells were washed in cold PBS, pelleted again by centrifugation, and then resuspended to a final concentration of 5-10 x10^6^ cells/ml in PBS supplemented with 1% BSA. Cell suspensions were filtered into round-bottomed tubes with cell-strainer caps (352235; Falcon). DR ESCs were sorted using an LSR II flow cytometer (BD Biosciences) and CRISPR-Cas9 edited E14 ESCs were sorted using FACSAria III flow cytometer (BD Biosciences), and analysis was performed using FACSDiva software (BD Biosciences). Statistical analysis was performed in Excel (Microsoft) using two-tailed t-test.

### RNA extraction, cDNA synthesis, and qPCR

Total RNA was isolated from ESCs at the indicated times by using TRIzol Reagent (15596026; Ambion) according to the manufacturer’s protocol with the following modifications: after washing cells with PBS 800 μl TRIzol was added to cells in a six-well plate and the pellet was rinsed in 75% EtOH. RNA purity was assessed on a Nanodrop spectrometer. cDNA was synthesized using the iScript cDNA Synthesis Kit according to manufacturer’s protocol (170-8891; Bio-Rad Laboratories). Quantitative PCR was performed with iQ SYBR® Green Supermix (170-8880; Bio-Rad Laboratories) according to the manufacturer’s protocol on a QuantStudio 6 Flex Real-Time PCR System (Applied Biosystems), with data analyzed using GraphPad Prism 6 software. For the stem cell lineage plate array, RNA was collected using TRIzol as indicated above. cDNA was synthesized using the RT2 First Strand Synthesis Kit according to manufacturer’s protocol (33041; Qiagen) qPCR primer sequences included:

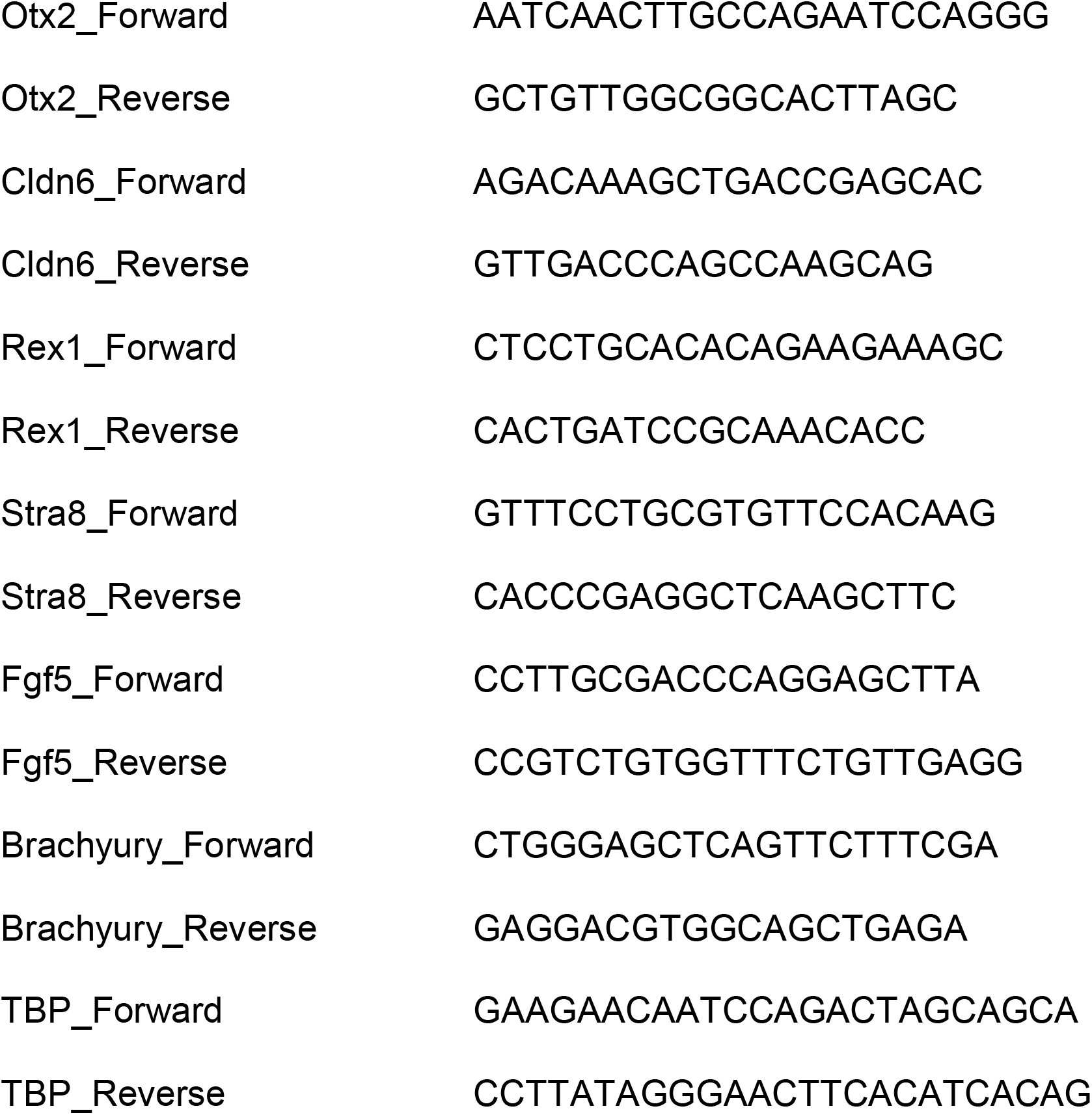

### Immunoblotting

Cells were lysed for 10 min in RIPA buffer (2.5 mM HEPES pH 7.5, 150 mM NaCl, 3mM KCl, 1% NP-40, 0.5% deoxycholate, 0.1% SDS, 1 mM vanadate, and 5 mM NaF supplemented with protease and phosphatase inhibitors). Lysates were centrifuged at 13,000 rpm for 15 min to obtain a post-nuclear supernatant. Proteins were separated by SDS-PAGE and transferred onto Immobilon-P® PVDF transfer membranes (IPVH00010; EMD Millipore) as previously described (Haynes et al., 2011; Rana et al., 2015). Membranes were blocked with 5% non-fat milk in TBS containing 0.1% Tween (TBST) and incubated with primary antibodies overnight at 4°C. Primary antibodies included α-tubulin (1:2000; GT114; GeneTex), Phospho-Myosin Light Chain 2 Thr18/Ser19 E2J8F (1:1000; #95777; Cell Signaling Technology), ERK 1 C-16 (1:1000; sc-93; Santa Cruz Biotechnology, Inc.), Phospho-p44/42 MAPK Erk1/2 Thr202/Tyr204 (1:1000; #9101; Cell Signaling Technology), Grhl2 (1:1000; HPA004820; Sigma), Arp2 (1:1000; A6104; Sigma), and Arpc2 (1:1000; 07-227; EMD Millipore). After washing, membranes were incubated in TBST with 5% non-fat milk and horseradish peroxidase (HRP)-conjugated secondary antibodies (1:10,000; 170-6516 and 172-1019; Bio Rad Laboratories) for 1 h at room temperature. After washing, immunoreactivity was developed with enhanced femto chemiluminescence (1859022 and 1859023; Thermo Scientific) and imaged using a BioRad Chemidoc XRS. ImageJ software was used for semi-quantitative densitometry analysis. Data presentation and statistical analysis were preformed using Excel Analyze-it and GraphPad Prism 6 software.

### Library preparation and RNA sequencing

RNA was extracted with the RNeasy Mini kit (Qiagen, 74104) according to the manufacturer’s instructions and sample concentrations were determined by NanoDrop. RNA degradation and contamination were monitored on 1% agarose gels, RNA purity was checked using the NanoPhotometer spectrophotometer (IMPLEN, CA, USA), and RNA integrity and quantitation were assessed using the RNA Nano 6000 Assay Kit of the Bioanalyzer 2100 system (Agilent Technologies, CA, USA). A total of 9 RNA libraries were prepared with three paired biological replicates for each condition including control +LIF2i, control 72h -LIF2i, and CK666 72h -LIF2i. A total amount of 1 μg RNA per sample was used as input material for the RNA sample preparations. Sequencing libraries were generated using NEBNext Ultra RNA Library Prep Kit for Illumina (NEB, USA) following manufacturer’s recommendations and index codes were added to attribute sequences to each sample. Briefly, mRNA was purified from total RNA using poly-T oligo-attached magnetic beads. Fragmentation was carried out using divalent cations under elevated temperature in NEBNext First Strand Synthesis Reaction Buffer (5X). First strand cDNA was synthesized using random hexamer primer and M-MuLV Reverse Transcriptase (RNase H). Second strand cDNA synthesis was subsequently performed using DNA Polymerase I and RNase H. Remaining overhangs were converted into blunt ends via exonuclease/polymerase activities. After adenylation of 3’ ends of DNA fragments, NEBNext Adaptor with hairpin loop structure was ligated to prepare for hybridization. To select cDNA fragments of preferentially 150∼200 bp in length, the library fragments were purified with AMPure XP system (Beckman Coulter, Beverly, USA). Then 3 μl USER Enzyme (NEB, USA) was used with size-selected, adaptor-ligated cDNA at 37°C for 15 min followed by 5 min at 95°C before PCR. Then PCR was performed with Phusion High-Fidelity DNA polymerase, Universal PCR primers and Index (X) Primer. At last, PCR products were purified (AMPure XP system) and library quality was assessed on the Agilent Bioanalyzer 2100 system. The clustering of the index-coded samples was performed on a cBot Cluster Generation System using PE Cluster Kit cBot-HS (Illumina) according to the manufacturer’s instructions. After cluster generation, the library preparations were sequenced on an Illumina platform and paired-end reads were generated with >20 million reads per sample. The above protocol, with the exception of RNA extraction, was performed externally by Novogene Co. Ltd (USA).

### RNA sequencing analysis

Quality assessment and basic processing of the reads was performed using the FastQC program (http://www.bioinformatics.babraham.ac.uk/projects/fastqc). Sequencing adapters were trimmed from the 3′ ends of the reads using cutadapt (v.1.8.1; https://pypi.python.org/pypi/cutadapt/1.8.1). We quantified transcript abundance with Kallisto (Bray et al., 2016) and built index with reference to the GRCh38 reference transcriptome. Expression analysis was performed using Sleuth (Pimentel et al., 2017) to assess differentially expressed genes between +LIF2i, control -LIF2i, and CK666 -LIF2i. Differentially expressed genes were identified using the Wald test with a cut-off of qval <0.05. Gene ontology enrichment analysis performed using Enrichr (Chen et al., 2013; Kuleshov et al., 2016). Heatmap figures were generated with Python using pandas dataframe (McKinney, 2010) input to the seaborn library (Waskom, 2021) in matplotlib (Hunter, 2007).

### Dataset acquisition

CHIP-seq data of TBX3 binding in mESCs was available from NCBI (GEO Series accession number: GSE19219) (Han et al., 2010). Microarray data from shRNA TBX3 knockdown mESCs was available from NCBI (GEO Series accession number: GSE26520) (Nishiyama et al., 2013). RNA-seq data from TBX3-HI and TBX3-LO mESCs was available from NCBI (GEO Series accession number: GSE73862) (Russell et al., 2015). CHIP-seq data of MRTF binding in NIH3T3 fibroblasts was available from NCBI (GEO Series accession number: GSE45888) (Esnault et al., 2014).

## Data and code availability

RNA-sequencing data generated during this study have been deposited in Gene Expression Omnibus (https://www.ncbi.nlm.nih.gov/geo/) under Accession code GEO: GSE175391. Software/packages used to analyze the dataset are freely available.

## Acknowledgments

We thank A. Smith (University of Cambridge) for providing the E14 cell line and R. Blelloch (UCSF) for providing V6.5 wild type and DR cells. We also thank T. Nystul (UCSF), M. Welch (UC Berkeley), and the Barber Lab for helpful discussions. This work was supported by NIH grants GM116384 and CA197855 to DLB. FMA was supported by an HHMI Gilliam Fellowship, a UCSF Moritz-Heyman Discovery Fellowship, and an NIGMS T32GM008568 training grant.

## Author contributions

DLB conceived the hypothesis, which was developed by FMA. DLB and FMA obtained and analyzed data as well as assembled figures and wrote the manuscript.

The authors declare no competing interests.

## Figure Legends

Supplemental Video 1

**E14 colony morphology in the presence of LIF2i**. DIC time-lapse recording of E14 cells with images taken every 10 min for 18h on gelatin-coated glass, 40x.

Supplemental Video 2

**E14 colony morphology at 24-42h -LIF2i**. DIC time-lapse recording of E14 cells with images taken every 10min for 18h on gelatin-coated glass, 40x.

**Supplemental Figure 1.**
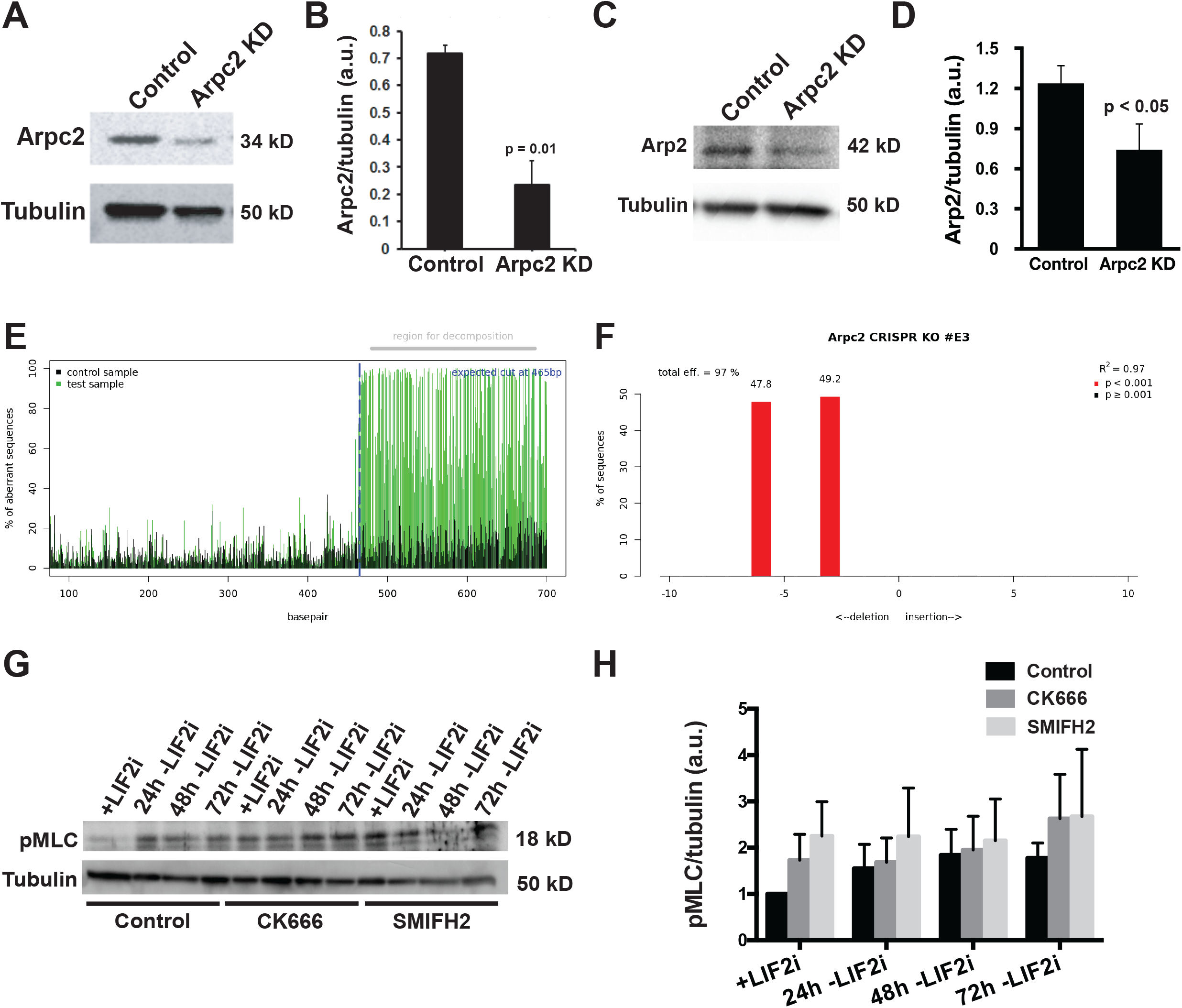
**(A)** Representative immunoblot of lysates from E14 control or Arpc2 KD mESCs probed for ARPC2 or tubulin as a loading control. **(B)** Semiquantitative densitometry of immunoblots described in (A), with data showing means ± SEM of 3 independent cell preparations. **(C)** Representative immunoblot of lysates from E14 control or Arpc2 KD mESCs probed for ARP2 or tubulin as a loading control. **(D)** Semiquantitative densitometry of immunoblots described in (C), with data showing means ± SEM of 3 independent cell preparations. **(E)** Profile of CRISPR-Cas9 edited E14 mESCs using TIDE webtool (https://tide.deskgen.com) to quantify editing efficacy by sequence aberration compared to control cells and to **(F)** identify the predominant indel (Brinkman *et al*., Nucleic Acids Reserch 2014). **(G)** Representative immunoblot of lysates from E14 mESCs over 72h timecourse -LIF2i in the absence (control) or presence of CK666 or SMIFH2 probed for pMLC and tubulin as a loading control. **(H)** Semiquantitative densitometry of immunoblots described in (G), with data showing means ± SEM of 3 independent cell preparations. Data were analyzed by two tailed unpaired Student’s *t*-test with a significance level of *p*<0.05.

**Supplemental Figure 2.**
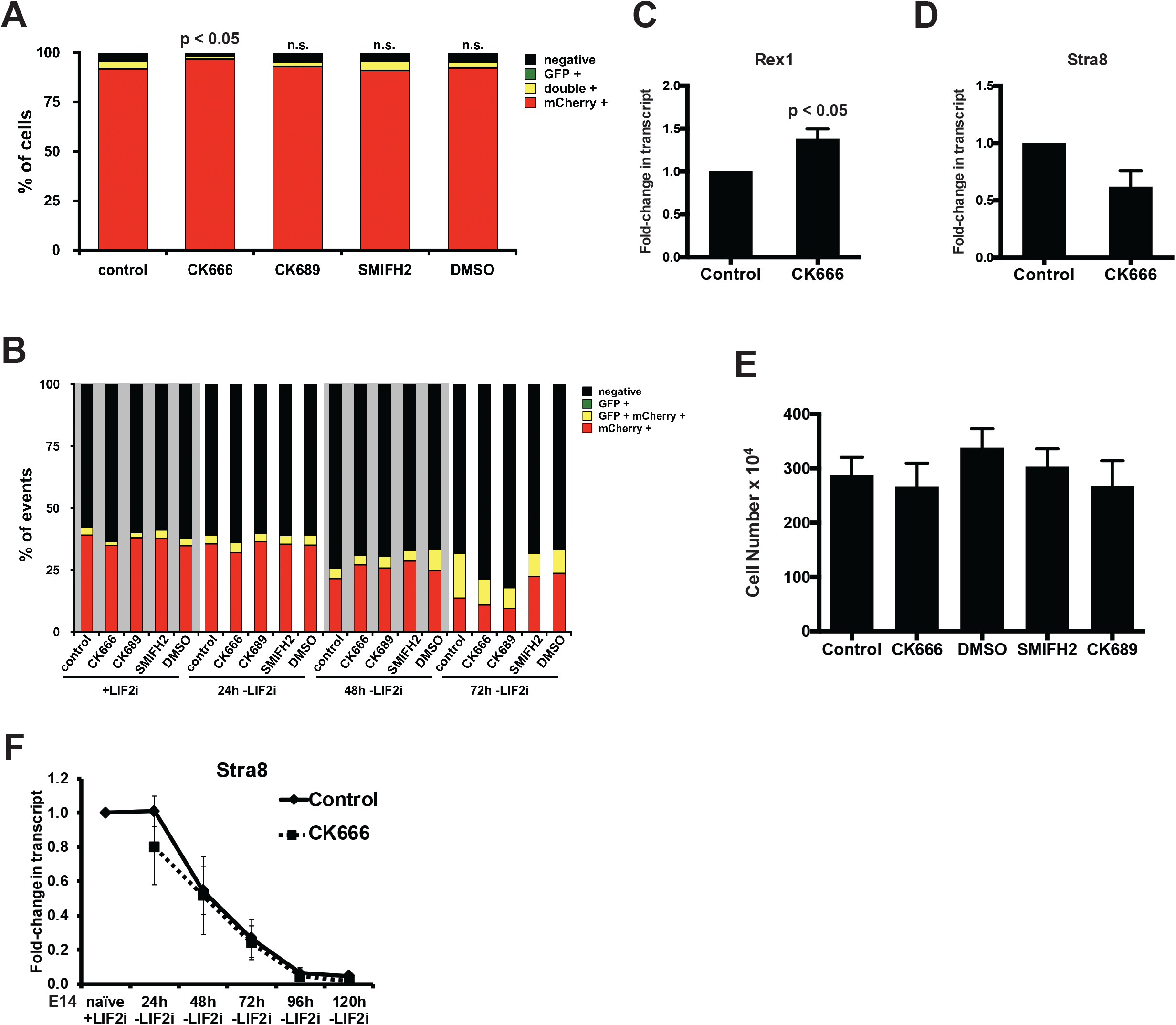
**(A)** FACS of V6.5 DR mESCs at 72h with LIF2i in the absence (control) or presence of CK666, CK689 inactive analog of CK666, SMIFH2, or DMSO vehicle, with indicated data from 6 independent cell preparations. **(B)** FACS of V6. 5 DR mESCs stained with DAPI during 72h - LIF2i timecourse in the absence (control) or presence of CK666, CK689 inactive analog of CK666, SMIFH2, or DMSO vehicle, with indicated data from 6 independent cell preparations to identify the differentiation status of only dead or dying cells. **(C)** RT-qPCR for *Rex1* and **(D)** *Stra8* in E14 mESCs at 120h with LIF2i in the absence (control) or presence of CK666 with indicated data showing means ± SEM of 4 independent cell preparations normalized to *TBP*. **(E)** Number of E14 mESCs after 72h -LIF2i in the absence (control) or presence of CK666, CK689 inactive analog of CK666, SMIFH2, or DMSO vehicle, with indicated data showing means ± SEM of 3 independent cell preparations. **(F)** RT-qPCR for *Stra8* in E14 mESCs during 120h - LIF2i timecourse in the absence (control) or presence of CK666, with indicated data showing means ± SEM of 4 independent cell preparations normalized to *TBP*. Data were analyzed by two tailed unpaired Student’s *t*-test with a significance level of *p*<0.05.

**Supplemental Figure 3.**
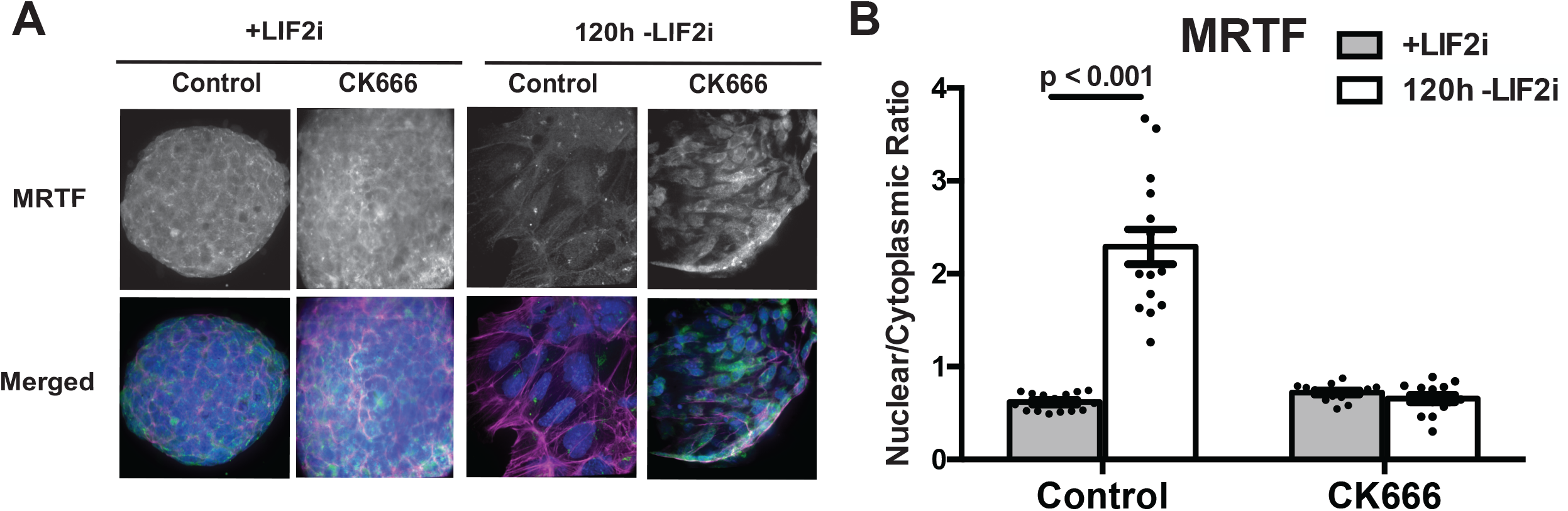
**(A)** Confocal images of E14 mESCs +LIF2i and at 120h -LIF2i in the absence or presence of CK666 immunolabeled for MRTF (green) and stained for F-actin with rhodamine phalloidin (magenta) and for nuclei with DAPI (blue). **(B)** Quantified nuclear to cytoplasmic ratio of MRTF immunolabeling shown in (A) indicating means ± SEM of 3 independent cell preparations. Data were analyzed by two tailed unpaired Student’s *t*-test with a significance level of *p*<0.05.

